# Next-generation sequencing-based bulked segregant analysis without sequencing the parental genomes

**DOI:** 10.1101/2021.02.08.430275

**Authors:** Jianbo Zhang, Dilip R. Panthee

## Abstract

The genomic region(s) that controls a trait of interest can be rapidly identified using BSA-Seq, a technology in which next-generation sequencing (NGS) is applied to bulked segregant analysis (BSA). We recently developed the significant structural variant method for BSA-Seq data analysis that exhibits higher detection power than standard BSA-Seq analysis methods. Our original algorithm was developed to analyze BSA-Seq data in which genome sequences of one parent served as the reference sequences in genotype calling, and thus required the availability of high-quality assembled parental genome sequences. Here we modified the original script to allow for the effective detection of the genomic region-trait associations using only bulk genome sequences. We analyzed a public BSA-Seq dataset using our modified method and the standard allele frequency and G-statistic methods with and without the aid of the parental genome sequences. Our results demonstrate that the genomic region(s) associated with the trait of interest could be reliably identified only via the significant structural variant method without using the parental genome sequences.

**Significance Statement:** BSA-Seq can be utilized to rapidly identify structural varianttrait associations, and our modified significant structural variant method allows the detection of such associations without sequencing the parental genomes, leading to further lower the sequencing cost and making BSA-Seq more accessible to the research community and more applicable to the species with a large genome.

---

Bulked segregant analysis (BSA) was developed for the quick identification of genetic markers associated with a trait of interest (1, 2). For a particular trait, two groups of individuals with contrasting phenotypes are selected from a segregating population. Equal amounts of DNA are pooled from each individual within a group. The pooled DNA samples are then subjected to analysis, such as restriction fragment length polymorphism (RFLP) or random amplification of polymorphic DNA (RAPD). Fragments unique to either group are potential genetic markers that may link to the gene(s) that control phenotypic expression for the trait of interest. Candidate markers are further tested against the population to verify the marker-trait associations. With the recent dramatic reductions in cost, next-generation sequencing (NGS) has been applied to more and more BSA studies (3–7). This new technology is referred to as BSA-Seq. In BSA-Seq, pooled DNA samples are not subjected to RFLP/RAPD analysis, but are directly sequenced instead. Genome-wide structural variants between bulks, such as single nucleotide polymorphisms (SNP) and small insertions/deletions (InDel), are identified based on the sequencing data. Genomic regions linked to the trait-controlling gene(s) are then identified based on the enrichment of the SNP/InDel alleles in those regions in each bulk. The time-consuming and labor-intensive marker development and genetic mapping steps are eliminated in the BSA-Seq method. Moreover, SNPs/InDels can be detected genome-wide via NGS, which allows for the reliable identification of trait-associated genomic regions across the entire genome.

For each SNP/InDel in a BSA-Seq dataset, the base (or oligo in the case of an InDel) that is the same as in the reference genome is termed the reference base (REF), and the other base is termed the alternative base (ALT). Because each bulk contains many individuals, the vast majority of SNP loci in the dataset have both REF and ALT bases. For each SNP, the number of reads of its REF/ALT alleles is termed allele depth (AD). Because of the phenotypic selection via bulking, for trait-associated SNPs, the ALT allele should be enriched in one bulk while the REF allele should be enriched in the other. However, for SNPs not associated with the trait, both ALT and REF alleles would be randomly segregated in both bulks, and neither enriched in either bulk. Hence these four AD values can be used to assess how likely a SNP/InDel is associated with the trait.

We have previously developed the significant structural variant method for BSA-Seq data analysis (8). In this method, a SNP/InDel is assessed with Fisher’s exact test using the AD values of both bulks. A SNP/InDel is considered significant if the P-value of Fisher’s exact test is lower than a specific cut-off value, e.g., 0.01. A genomic region normally contains many SNPs/InDels. The ratio of the significant structural variants to the total structural variants is used to judge if this genomic region is associated with the trait of interest. We tested this method using the BSA-Seq data of a rice cold-tolerance study (9). One of the parents in this study was rice cultivar *Oryza sativa* ssp. *japonica* cv. Nipponbare. Its high-quality assembled genome sequences were used as the reference sequences for SNP/InDel calling as well, which makes the genotype calling and SNP/InDel filtering very straightforward: any locus in any bulk that is different from the REF allele is a valid SNP/InDel (8).

Only high-quality assembled genome sequences can serve as the reference sequences in genotype calling, an essential step in BSA-Seq data analysis. For most species, however, such sequences are available for only a single or limited number of lines. If lines without high-quality assembled genome sequences are used as the parents in BSA-Seq studies, the parental genomes are often sequenced via NGS for the determination of the parental origin of SNP alleles and the identification of parental heterozygous SNPs. Modification of our original method to allow the analysis of BSA-Seq data in the absence of assembled or NGS-generated parental genome sequences would provide greater flexibility and significantly reduce sequencing costs. Hence, we modified our original script to allow for the identification of the false-positive SNPs/InDels and part of the heterozygous loci in the parents without the aid of the parental genome sequences. Using the modified script, along with the scripts for the standard G-statistic and allele frequency methods (10, 11), we analyzed a public BSA-Seq dataset using either the genome sequences of both the parents and the bulks, or the bulk genome sequences alone. The results revealed that reliable detection of genomic region-trait associations can be achieved only via our modified script when using only the bulk genome sequences.

## Materials and Methods

The sequencing data used in this study were generated by Lahari *et al.* (12). Using the allele frequency method, the authors identified a single locus for root-knot nematode resistance in rice. In that study, the parents of the F2 population were LD24 and VialoneNano, yielding an F2 population size of 178 (plants), and both the resistant bulk and the susceptible bulk contained 23 plants each. The DNA samples of both the parents and the bulks were sequenced using Illumina MiSeq Sequencing System and MiSeq v3 chemistry.

The BSA-Seq sequencing data (ERR2696318: parent LD24; ERR2696319: parent VialoneNano; ERR2696321: the resistant bulk from the F2 population; ERR2696322: the susceptible bulk from the F2 population) were downloaded from the European Nucleotide Archive (ENA) using the Linux program wget, and the rice reference sequence (Release 47) was downloaded from https://plants.ensembl.org/Oryza_sativa/Info/Index. Sequencing data preprocessing and SNP calling were performed as described previously (8). When analyzing the BSA-Seq data with the genome sequences of both the parents and the bulks, bulk/parent SNP calling was performed separately. The common SNPs of the two SNP datasets were used for the downstream analysis.

The SNP dataset generated via SNP calling was processed with our Python script to identify significant SNP-trait associations. A single script containing all the three methods is available on the website https://github.com/dblhlx/PyBSASeq. The workflow of the scripts is as follows:

1. Read the.tsv input file generated via SNP calling into a Pandas DataFrame.
2. Perform SNP filtering on the Pandas DataFrame.
3. Identify the significant SNPs (sSNPs) via Fisher’s exact test (the significant structural variant method), calculate the ΔAF (allele frequency difference between bulks) values (the allele frequency method), or calculate the G-statistic values (the G-statistic method) using the four AD values (AD_ref1_ and AD_alt1_ of bulk 1 and AD_ref2_ and AD_alt2_ of bulk 2) of each SNP in the filtered Pandas DataFrame.
4. Use the sliding window algorithm to plot the sSNP/totalSNP ratios, the ΔAF values, or the G-statistic values against their genomic positions.
5. Estimate the threshold of the sSNP/totalSNP ratio, the ΔAF, or the G-statistic via simulation. The thresholds are used to identify the significant peaks/valleys in the plots generated in step 4.

Identification of the sSNPs, calculation of the sSNP/totalSNP ratios, the G-statistic values, or the ΔAF values, and estimation of their thresholds were carried out as described previously (8). The 99.5^th^ percentile of 10000 simulated sSNP/totalSNP ratios or G-statistic values was used as the threshold for the significant structural variant method or the G-statistic method, and the 99% confidence interval of 10000 simulated ΔAF values was used as the threshold for the allele frequency method. For all methods, the size of the sliding windows is 2Mb and the incremental step is 10kb. In our previous work, a parent was the japonica rice cultivar nipponbare, and its genome sequences were used as the reference sequences for SNP/InDel calling. In the current dataset, the parents were LD24 and VialoneNano; many false-positive SNPs/InDels and heterozygous loci in the parents would be included in the dataset if analyzing the BSA-Seq data using the original script. Hence, SNP filtering is carried out a little differently from previously described (8), and its details are below (see Table S1 for examples):

- Unmapped SNPs or SNPs mapped to the mitochondrial or chloroplast genome
- SNPs with an ‘NA’ value in any column of the DataFrame
- SNPs with zero REF read and a single ALT allele in both bulks/parents
- SNPs with three or more ALT alleles in any bulk/parent
- SNPs with two ALT alleles and its REF read is not zero in any bulk/parent
- SNPs in which the bulk/parent genotypes do not agree with the REF/ALT bases
- SNPs in which the bulk/parent genotypes are not consistent with the AD values
- SNPs with a genotype quality (GQ) score less than 20 in any bulk
- SNPs with very high reads
- SNPs heterozygous in any parent when parental genome sequences are available

Additionally, for SNPs with two ALT alleles and zero REF read in both bulks/parents, the REF allele is replaced with the first allele in the ‘ALT’ field, its ALT allele is replaced with the second allele in the original ‘ALT’ field. The REF read, and a comma after it, are removed from both the allele depth (AD) fields (one for each bulk/parent). This step is carried out before checking the genotype agreement between bulks and the REF/ALT fields. When parental genome sequences are involved, the common SNP set is identified before filtering out the SNPs with a low GQ score in the parental SNP dataset.

The tightly linked SNP alleles from the same parent tend to segregate together and should have a similar extent of allele enrichment, and thus similar AD values. In a SNP dataset, the genotypes of each bulk/parent are represented as ‘GT_ref_/GT_alt_ when a SNP contains both the REF base and the ALT base in the genotype (GT) field, and the AD values in each bulk/parent is represented as ‘AD_ref_,AD_alt_’. The genotype and the AD value of the REF allele are always placed first in both fields. For a SNP locus in the.tsv input file, the allele having the same genotype as that in the reference genome is defined as the REF allele. However, it is highly unlikely that all of the SNP alleles in a parent are the same as those in the reference genome, except in instances where reference genome sequences used in SNP calling are from one of the parents as in the case of the cold-tolerance study as mentioned above (9). It is necessary to place the genotypes and AD values of all SNP alleles from one parent (e.g., LD24) in the REF position, and those from the other parent (e.g., VialoneNano) to the ALT position in the GT and AD fields to make the bulk dataset consistent. Thus, for a particular SNP, if the REF base in the.tsv file is different from the genotype of LD24 (either parent will work), its GT/AD values would be swapped, e.g., ‘G/A’ to ‘A/G’ and ‘19,9’ to ‘9,19’. AD/GT swapping is performed following SNP filtering and is performed only when the parental genome sequences are used to aid BSA-Seq data analysis. Equation 1 is used for ΔAF calculation. AD swapping ensures that adjacent SNPs have similar ΔAF values.

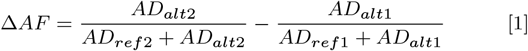

## Results

The original sequence reads were 3.9G, 3.8G, 3.4G, and 3.5G; they became 3.8G, 3.6G, 3.3G, and 3.4G after quality control, respectively, in ERR2696318 (parent LD24), ERR2696319 (parent VialoneNano), ERR2696321 (the resistant bulk), and ERR2696322 (the susceptible bulk), which correspond to 8.8×, 8.5×, 7.6×, and 7.9× coverage, respectively (12). The preprocessed sequences were used for SNP calling to generate a SNP dataset, which was analyzed using the modified significant structural variant method, the G-statistic method, and the allele frequency method with or without the aid of the parental genome sequences.

### BSA-Seq data analysis using the genome sequences of both the parents and the bulks

The SNP calling-generated parent/bulk SNP dataset was processed with the Python script PyBSASeq_WP.py. SNP filtering was performed as described in the Materials and Methods section. The parental SNP dataset was processed first, and the SNPs heterozygous in any parent were eliminated because all algorithms assume all SNP loci are homozygous in the parental lines. Threshold estimation is based on this assumption. Although most rice breeding lines should be homozygous in most loci, more than 7% heterozygous SNP loci (2011062 homozygous and 153000 heterozygous) were identified in the parental SNP dataset. However, the GATK’s variant calling tools are designed to be very lenient in order to achieve a high degree of sensitivity (https://gatk.broadinstitute.org/hc/en-us/articles/360035535932-Germline-short-variant-discovery-SNPs-Indels-), we cannot rule out the possibility that some of the heterozygous loci were caused by sequencing artifacts. The bulk SNP dataset was processed second. The SNPs with the same chromosome ID and the same genomic coordinate in both datasets were considered common SNPs. Common SNPs in the bulk dataset were used to detect SNP-trait associations for all three methods.

#### The significant structural variant method

Each SNP in the dataset was tested via Fisher’s exact test using its four AD values, and SNPs with P-values less than 0.01 were defined as sSNPs. The chromosomal distributions of the sSNPs and the total SNPs are summarized in Table 1. Using the sliding window algorithm, the genomic distribution of the sSNPs, the total SNPs, and the sSNP/totalSNP ratios of sliding windows were plotted against their genomic position (Figure 1a and Figure 1b). A genome-wide threshold was estimated as 0.0538 via simulation as described previously (8). Two peaks above the threshold were identified: a minor one on chromosome 9 and a major one on chromosome 11. The position of the peak on chromosome 9 was at 1.11Mb, the sliding window contained 230 sSNPs and 3738 total SNPs, corresponding to an sSNP/totalSNP ratio of 0.0615; the position of the peak on chromosome 11 was at 26.44 Mb, the sliding window contained 675 sSNPs and 1139 total SNPs, corresponding to an sSNP/totalSNP ratio of 0.5926. The sliding window-specific threshold was estimated for each peak via simulation, and the values were 0.0551 and 0.0623, respectively, indicating both peaks were significant. Both values are higher than the genome-wide threshold, probably due to the lower amounts of total SNPs in these sliding windows. The average SNPs per sliding window was 5893.

**Figure 1.**
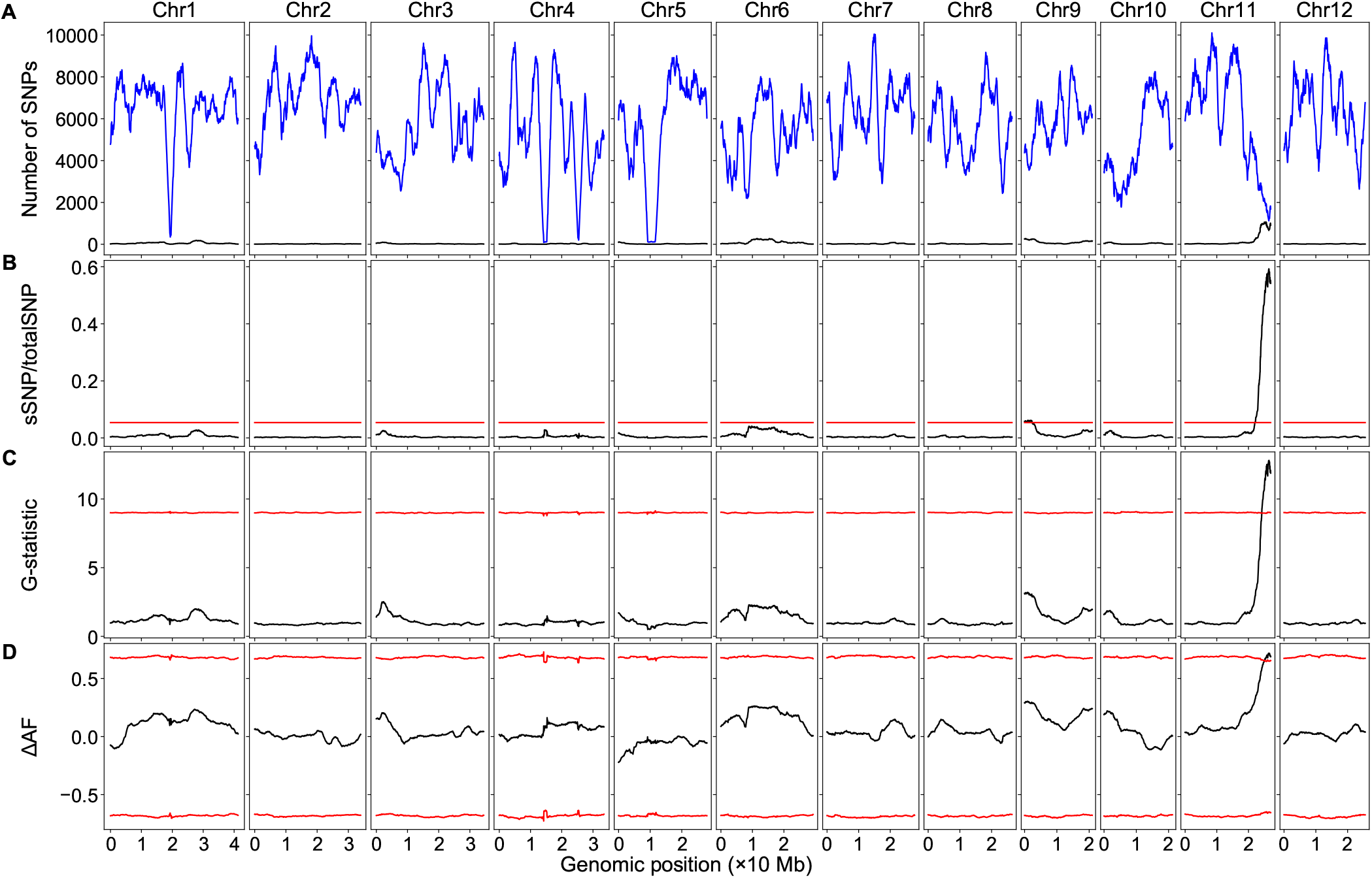
BSA-Seq data analysis using the genome sequences of both the parents and the bulks. The red lines/curves are the thresholds. **(A)** Genomic distributions of sSNPs (blue) and totalSNPs (black). **(B)** Genomic distributions of sSNP/totalSNP ratios. **(C)** Genomic distributions of G-statistic values. **(D)** Genomic distributions of ΔAF values.

**Table 1.**
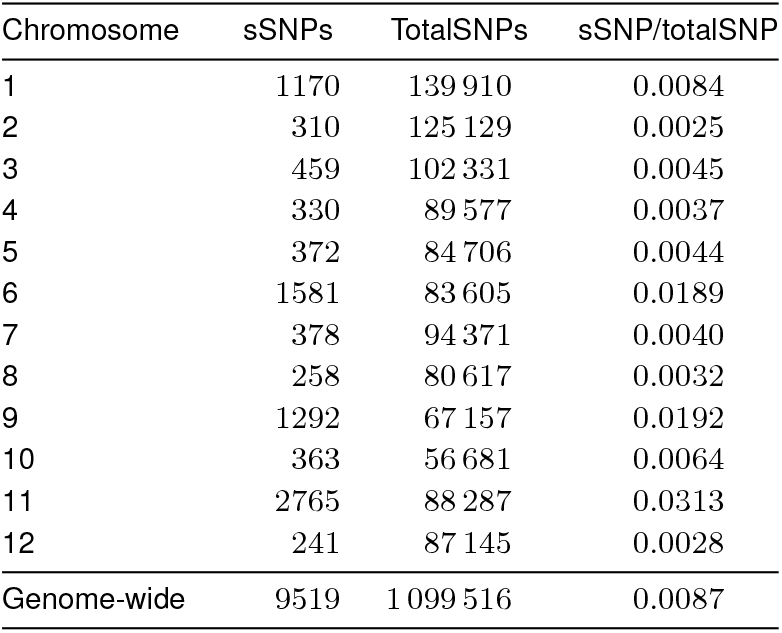
Chromosomal distribution of SNPs - using the genome sequences of both the parents and the bulks.

| Chromosome | sSNPs | TotalSNPs | sSNP/totalSNP |
| --- | --- | --- | --- |
| 1 | 1170 | 139 910 | 0.0084 |
| 2 | 310 | 125 129 | 0.0025 |
| 3 | 459 | 102 331 | 0.0045 |
| 4 | 330 | 89 577 | 0.0037 |
| 5 | 372 | 84 706 | 0.0044 |
| 6 | 1581 | 83 605 | 0.0189 |
| 7 | 378 | 94 371 | 0.0040 |
| 8 | 258 | 80 617 | 0.0032 |
| 9 | 1292 | 67 157 | 0.0192 |
| 10 | 363 | 56 681 | 0.0064 |
| 11 | 2765 | 88 287 | 0.0313 |
| 12 | 241 | 87 145 | 0.0028 |
| Genome-wide | 9519 | 1 099 516 | 0.0087 |

#### The G-statistic method

The G-statistic value of each SNP in the dataset was calculated, and its threshold was estimated via simulation as described previously (8). Using the sliding window algorithm, the G-statistic value of each sliding window, the average G-statistic values of all SNPs in that sliding window, was plotted against its genomic position (Figure 1c), and the curve pattern was very similar to that in Figure 1b. A significant peak was identified on chromosome 11; its position was at 26.49 Mb, its G-statistic value was 12.8120, well above the threshold 9.0224 (99.5^th^ percentile).

#### The allele frequency method

The ΔAF value of each SNP in the dataset was calculated, and the ΔAF threshold of the SNP was estimated via simulation as described previously (8). Using the sliding window algorithm, the ΔAF value of each sliding window, the average ΔAF values of all SNPs in that sliding window, was plotted against its genomic position (Figure 1d). A significant peak on chromosome 11 was identified, the peak position was located at 26.45 Mb, its ΔAF value was 0.7173, and the 99% confidence interval was −0.6508 to 0.6497.

### BSA-Seq data analysis using only the bulk genome se-quences

The SNP calling-generated bulk SNP dataset was processed with the Python script PyBSASeq.py. All the methods and parameters were the same as above; the only difference was that the parental SNP dataset was not used.

#### The significant structural variant method

The chromosomal distribution of the sSNPs and total SNPs are summarized in Table 2. The total number of SNPs was 1 346 185 here, much higher than the above, which was 1 099 516. The ge-nomic distribution of the sSNPs, the total SNPs, and the sSNP/totalSNP ratios of the sliding windows are presented in Figure 2a and Figure 2b. The patterns of the curves were very similar to those in Figure 1a and Figure 1b. One of the obvious differences was that sSNP/totalSNP ratios of the sliding windows were much lower than those in Figure 1b, leading to missing the minor locus on chromosome 9. Only the peak on chromosome 11 was significant; it was located at 26.96 Mb, a 520 kb shift compared to Figure 1b. The sliding window contained 1122 sSNPs and 2945 total SNPs, corresponding to a 0.3810 sSNP/totalSNP ratio, well above the genome-wide threshold (0.0535) and the sliding window specific threshold (0.0601). The average SNPs per sliding window was 7215.

**Figure 2.**
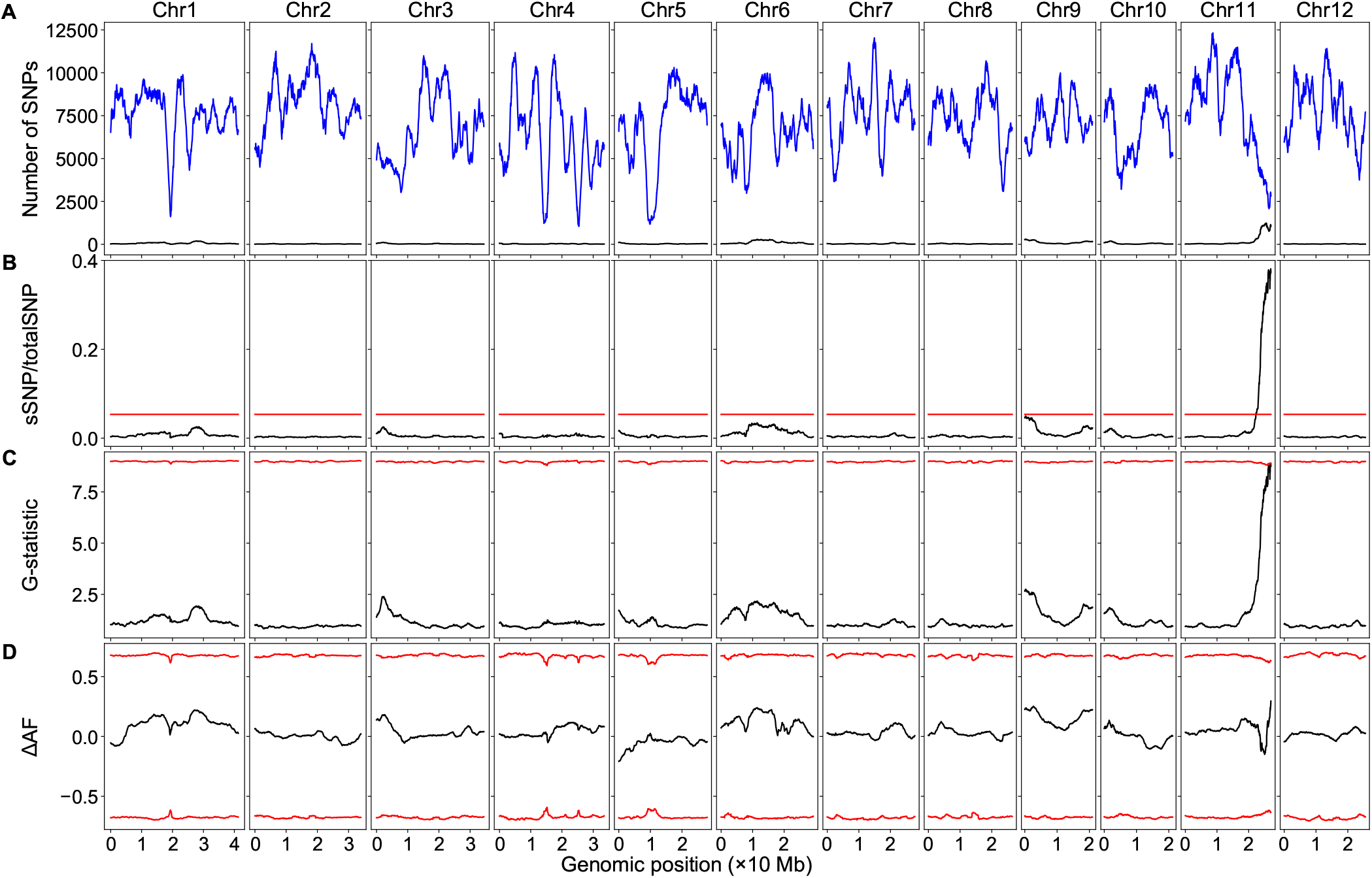
BSA-Seq data analysis using only the bulk genome sequences. The red lines/curves are the thresholds. **(A)** Genomic distributions of sSNPs (blue) and totalSNPs (black). **(B)** Genomic distributions of sSNP/totalSNP ratios. **(C)** Genomic distributions of G-statistic values. **(D)** Genomic distributions of ΔAF values.

**Table 2.** Chromosomal distribution of SNPs - using only the bulk genome sequences.

| Chromosome | sSNPs | TotalSNPs | sSNP/totalSNP |
| --- | --- | --- | --- |
| 1 | 1335 | 163 260 | 0.0082 |
| 2 | 391 | 146 877 | 0.0027 |
| 3 | 578 | 120 319 | 0.0048 |
| 4 | 442 | 110 952 | 0.0040 |
| 5 | 481 | 103 362 | 0.0047 |
| 6 | 1724 | 103 416 | 0.0167 |
| 7 | 459 | 114 564 | 0.0040 |
| 8 | 373 | 103 385 | 0.0036 |
| 9 | 1410 | 82 744 | 0.0170 |
| 10 | 572 | 78 206 | 0.0073 |
| 11 | 3120 | 112 719 | 0.0277 |
| 12 | 281 | 106 381 | 0.0026 |
| Genome-wide | 11 166 | 1 346 185 | 0.0083 |

#### The G-statistic method

The patterns of the G-statistic value plot (Figure 2c) were very similar to that in Figure 1c, but the G-statistic values were significantly lower than those in Figure 1c, and the threshold did not change much. Only a single sliding window was above the threshold (8.8953), its position was at 29.96 Mb, and its G-statistic value was 8.9060.

#### The allele frequency method

Without the aid of the parental genome sequences, the pattern of the ΔAF curve of chromosome 11 (Figure 2d), especially the genomic region associated with the trait, was drastically different from that in Figure 1d. Differences in the curve patterns were observed in other chromosomes as well, but they were relatively minor. All ΔAF values were within the 99% confidence interval, although AD swapping was performed on only 67 396SNPs, 6.1% of total SNPs.

## Discussion

We tested how parental genome sequences affected the detection of SNP-trait associations via BSA-Seq using a dataset of the rice root-knot nematode resistance. Using the genome sequences of both the parents and bulks, a major locus on chromosome 11 and a minor locus on chromosome 9 were detected via the significant structural variant method. However, only the major locus was detected via the G-statistic method and the allele frequency method. The positions of the peaks detected via different methods were not the same, but they were very close to each other. Using only the bulk genome sequences, the major locus can be detected via only the significant structural variant and G-statistic methods. The allele frequency method uses the ΔAF value of a SNP to measure allele (REF/ALT) enrichment in the SNP locus, and the G-statistic method uses the G-statistic value of a SNP to measure the allele enrichment; ΔAF and G-statistic are parameters at the SNP level, therefore, both methods use a SNP level parameter to identify significant sliding windows for the detection of the genomic region-trait associations. The significant structural variant method, however, uses the sSNP/totalSNP ratio, a parameter at the sliding window level, to measure the sSNP enrichment in a sliding window for the identification of the trait-associated genomic regions. A SNP normally has less than 100 reads because of the cost concern, while a sliding window normally contains thousands of SNPs. Thus, the significant structural variant method has much higher statistical power, which is consistent with our observation. Our results revealed that the parental genome sequences did not much affect the plot patterns of the sSNP/totalSNP ratios and the G-statistic values. However, the plot patterns of the ΔAF value of chromosome 11 were altered dramatically when the parental genome sequences were not used.

The significant structural variant method assesses if a SNP is likely associated with the trait via Fisher’s exact test. The greater the ALT proportion differences between the bulks, the less the P-value of the Fisher’s exact test, and the more likely the SNP is associated with the trait. Fisher’s exact test takes a numpy array or a Python list as its input, the same P-value will be obtained with either [[AD_ref1_, AD_alt1_], [AD_ref2_, AD_alt2_]] or [[AD_alt1_, AD_ref1_], [AD_alt2_, AD_ref2_]] as its input. The G-statistic method assesses if a SNP is likely associated with the trait via the G-test; the greater the G-statistic value of a SNP, the more likely it contributes to the trait phenotype (11). The G-statistic values are the same with either input [[AD_ref1_, AD_alt1_], [AD_ref2_, AD_alt2_]] or [[AD_alt1_, AD_ref1_], [AD_alt2_, AD_ref2_]]. The order of the AD values (REF/ALT reads) in bulks does not affect the P-value of Fisher’s exact test or the G-statistic value of G-test, which is why the parental genome sequences-guided AD swapping does not alter the curve patterns of both methods. Therefore, theoretically, parental genome sequences are not required to identify genomic region-trait associations in either the significant structural variant method or the G-statistic method.

When the parental genome sequences were used, AD value swapping was performed for the SNPs in which the genotype of LD24 was different from the REF base, and the ΔAF values of these SNPs were calculated based on the swapped AD values using equation 1. AD swapping makes the adjacent SNP alleles from the same parent have similar AD values and similar ΔAF values. The ΔAF values of such SNPs were calculated using equation 2 if not performing AD swapping. Equation 2 can be converted to equation 3, which produces an opposite value relative to that produced by equation 1. For two adjacent SNPs in LD24, where one SNP has the same genotype as the REF base while the other has the same genotype as the ALT base, they would have opposite ΔAF values if AD swapping is not performed. For the SNPs that do not contribute to the trait phenotype and are not linked to any trait-associated genomic regions, their ΔAF value should fluctuate around zero. The parental genome sequences will have less effect on the ΔAF value of the sliding windows containing such SNPs. However, for trait-associated SNPs, adjacent SNPs with opposite ΔAF values would cancel each other out and lower the ΔAF value of the sliding window significantly, which is the case observed on chromosome 11 in Figure 2d.

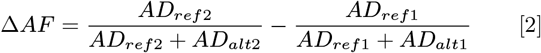

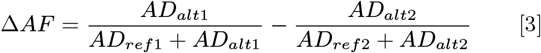

When the parental genome sequences were not used, the sSNP/totalSNP ratios and the G-statistic values were significantly lower. The peak sSNP/totalSNP ratio on chromosome 11 was 0.5926 in Figure 1b, while it was 0.3810 in Figure 2b; it was similar for the peak G-statistic values. The decreasing of sSNP/totalSNP ratio and the G-statistic value is likely caused by sequencing artifacts and heterozygosity in the parental lines. There were 1345185 SNPs in the bulk dataset when not using the parental genome sequences, while there were 1099516 SNPs in the dataset with the aid of the parental genome sequences. Comparison of the two SNP dataset revealed that 109445 SNPs were unique to the bulks. Because all the SNPs in the bulks are derived from the parental lines, crossing should not generate new SNPs; thus this category of SNPs was most likely caused by sequencing artifacts. The sequencing coverage in the bulk was less than eight, which is very low. Higher sequencing coverage would help decrease the number of SNPs derived from sequence artifacts. Additionally, 137 224 SNP were heterozygous in the parental lines. Without the parental genome sequences, this category of SNPs could not be filtered out from the bulk SNP dataset. However, these SNPs can be decreased via selfing the parental line more generations: five-generations selfing can decrease the heterozygosity of both parental lines to a maximum of 6.25%.

To determine how parental heterozygosity and sequencing artifacts affected the detection of genomic region-trait associations, we removed the heterozygous SNPs or the bulk-specific SNPs from the bulk SNP dataset, and analyzed the data separately. By removing the heterozygous SNPs, the peak on chromosome 11 was shifted to 26.28Mb for both the sSNP/totalSNP ratio and the G-statistic value, and the sSNP/totalSNP ratio of the peak was increased to 0.4835, well above the sliding window-specific threshold 0.0603. The G-statistic value of the peak was 10.8411, significantly higher than the threshold 8.9532 as well. By removing bulk-specific SNPs, the peak on chromosome 11 shifted to 26.49 Mb for both the sSNP/totalSNP ratio and the G-statistic value. The sSNP/totalSNP ratio of the peak and the sliding window-specific threshold were 0.4302 and 0.0637, respectively, and the G-statistic value of the peak and the threshold were 9.7591 and 8.9092, respectively. Although both the sSNP/totalSNP ratio and the G-statistic value were lower than above, they were still higher than their corresponding thresholds. While seemed the heterozygous SNPs affected the sSNP/totalSNP ratio and the G-statistic value a little more than the bulk-specific SNPs, it is more likely that both produced similar levels of noise for the sSNP/totalSNP ratio and the G-statistic value considering that the former was 27 779 greater than the latter. When using only the bulk genome sequences, the sSNP/totalSNP peak position on chromosome 11 was shifted 0.52 Mb (26.44 Mb to 26.96 Mb) due to the presence of the bulk-specific SNPs and the heterozygous SNPs in the dataset, but this is a very short distance for genetic mapping. Although only a single dataset was examined here, the genome-wide similarity of the sSNP/totalSNP curve patterns in Figure 1b and Figure 2b suggests that the significant structural method is highly reproducible using only the bulk genome sequences.

## Conclusions

The plotting pattern of the ΔAF values in the trait-associated genomic region was very different when using only the bulk genome sequences. Without the aid of the parental genome sequences, the ΔAF values of the sliding windows could not be correctly calculated; thus, the allele frequency method cannot be used to identify SNP-trait association. In contrast, the parental genome sequence does not affect the plotting patterns of both the significant structural variant method and the G-statistic method, but the sSNP/totalSNP ratios and the G-statistic values decreased significantly due to sequencing artifacts and/or heterozygosity of the parental lines. Because of its high detection power, major SNP-trait associations can still be reliably detected via the significant structural variant method even the sequence coverage was very low.

## Author contributions

JZ and DRP conceived the study. JZ developed the algorithm, wrote the Python code, analyzed the data, and wrote and edited the manuscript. DRP edited the manuscript and supervised the project.

The authors declare no conflict of interest.

## Acknowledgments

JZ was supported by the National Science Foundation grant [IOS-1546625 to DRP]. We are grateful to Lahari *et al.* for generating the sequencing data and making it available to the public. We thank Irene E. Palmer for critical review and thank Nathan Lynch for valuable comments. The manuscript was prepared using a modified version of the PNAS LATEX template.

